# Automated Locus Coeruleus segmentation and age-related signal changes across Neuromelanin-sensitive MRI and T1/T2-weighted images

**DOI:** 10.1101/2025.09.12.675918

**Authors:** Sun Hyung Kim, Martin Andreas Sytner, Joshua Neal, Ya-Yun Chen, Benjamin Katz, Tae-Ho Lee

**Affiliations:** Department of Psychiatry, University of North Carolina at Chapel Hill, USA; Department of Biomedical Engineering, University of Basel, Switzerland; Department of Psychology, Virginia Tech, USA; Department of Human Development and Family Science, Virginia Tech, USA; School of Neuroscience, Virginia Tech, USA

**Keywords:** Locus Coeruleus, Automatic segmentation, Developmental intensity change, Neuromelanin-sensitive MRI, T1-and T2-weighted MRI

## Abstract

The locus coeruleus (LC) plays a key role in stress response, attention, and cognitive function, with its integrity closely linked to neuromelanin accumulation. Neuromelanin-sensitive MRI (NM-MRI) has enabled in vivo imaging of the LC, but automated segmentation remains challenging due to its small size, low contrast, and anatomical variability. This study aimed to evaluate the accuracy of automatic LC segmentation using multi-atlas nonlinear warping applied to high-resolution T1/T2-weighted images and to compare LC intensity characteristics across NM-MRI, T1w, and T2w images. Additionally, we examined developmental changes in LC signal from childhood through adulthood. We analyzed MRI data from 134 participants (ages 8∼64) with NM-MRI, T1w, and T2w scans, alongside 652 participants (ages 6∼22) from the Human Connectome Project in Development (HCP-D). LC intensity measures were compared across imaging modalities and analyzed for age-related trends. Automatic LC segmentation achieved high spatial overlap with manual tracings (up to 95% after one voxel 2D-dilation). Statistically estimated high (or low)-intensity values (mean ± 2σ) from automated LC masks provided a more stable and noise-resilient alternative to local peak intensity measures. Age-related LC signal changes are most clearly observed in NM-MRI and T2w imaging, with significant developmental shifts emerging during adolescence. These findings support the conventional T2w imaging as an indirect marker of neuromelanin accumulation and highlight the possibility of age-specific analyses in LC imaging studies.

## Introduction

The locus coeruleus (LC) in the brainstem plays a central role of the stress response, attention regulation and other physiological and cognitive functions with dys-and modulating norepinephrine neurotransmitter level (Bell et al., 2023; Maness et al., 2022; Slavova et al., 2024). This LC has widespread projection throughout the brain, including the prefrontal cortex, amygdala/hippocampus, thalamus as well as the motor and sensory related cortex. Thus, LC dysfunction is not only one of potential mechanisms in emotional or degenerative issues but also intensively related the cortical structural or functional alteration.

The relationship between norepinephrine and neuromelanin in the LC is closely tied to the metabolic activity of the neurons (Wakamatsu et al., 2015). Neuromelanin is a dark pigment derived from the byproducts of catecholamine metabolism, specifically norepinephrine and dopamine. When norepinephrine is metabolized, some byproducts can lead to the formation of oxidized quinones. These oxidized compounds can polymerize into neuromelanin that has accumulated over time in neurons of LC. Neuromelanin accumulation is a natural result of the high metabolic activity of norepinephrine-producing neurons in the LC. The presence of neuromelanin in the LC indicates a history of norepinephrine synthesis and neuronal activity (Iannitelli & Weinshenker, 2023; Nikolenko et al., 2024; Sulzer et al., 2018; Wakamatsu et al., 2015). One of the most representative examples is in neurodegenerative diseases like Parkinson’s and Alzheimer’s where damaged LC neurons release neuromelanin, which can trigger inflammatory responses and oxidative stress, contributing to neuronal death. However, neuromelanin is not actively involved in neurotransmission but rather a byproduct of cellular metabolism (Capucciati et al., 2021; Moreno-García et al., 2021; Sulzer et al., 2018).

Neuromelanin-sensitive MRI (NM-MRI) has been used to detect the neuromelanin in LC. NM-MRI relies on the unique magnetic properties of neuromelanin, which binds to metals like iron and copper. These properties affect the local magnetic field, making neuromelanin paramagnetic and detectable using specialized MRI sequences. In degenerative neurological disorders, the NM-MRI has proven valuable in detecting structural changes associated with neuronal degeneration. Signal intensity reductions in the LC and substantia nigra pars compacta (SNc) have been observed, indicating a loss of neuromelanin-containing neurons. Notably, NM-MRI can detect signal alterations in the lateral SNc and LC in patients with Parkinson’s disease (PD), even during the early stages, and effectively differentiate PD patients from healthy individuals with high sensitivity and specificity (Ohtsuka et al., 2013; Wiesman et al., 2024). Emerging evidence suggests the potential application of NM-MRI in psychiatric disorders as well. Recent studies have provided the first in vivo demonstration of midbrain dopamine alterations in pediatric psychiatry, particularly in the obsessive-compulsive disorder and depression (Calarco et al., 2022; Pagliaccio et al., 2023). And the T2-star weighted MRI (T2*) is also sensitive for iron deposits, microbleeds, and susceptibility effects, making it useful for conditions like hemorrhage and neurodegeneration. The intensity value of T2* MRI in the LC may be the biological imaging indicators for the early diagnosis, severity, and follow-up evaluation of PD (Cao et al., 2023; Priovoulos et al., 2020).

Given the critical importance of the LC, it is necessary to define the region of interest (ROI) on at least one image with the highest quality among the multiple acquired scans (e.g. T1w/T2w or NM-MRI) to ensure accurate analysis. The LC is a cylindrical structure averaging 15mm in length and 2–2.5mm in diameter (Betts et al., 2019). Delineating this small region in NM-MRI has traditionally relied on conservative thresholding of a warped probabilistic atlas (Wiesman et al., 2024; Ye et al., 2021) or manual tracing (Neal et al., 2023; Ohtsuka et al., 2013). However, a single subject atlas approach has several limitations in brain tissue segmentation. Since it represents only one anatomical structure of single subject, it may have a risk to account for inter-individual variations in brain morphology, leading to segmentation errors. Additionally, as the atlas is derived from a specific population (e.g., an averaged template), it may not accurately capture anatomical differences due to age, pathology, or structural variability across individuals or specific group (Aljabar et al., 2009; Wang et al., 2014; Yaakub et al., 2020). Accurate segmentation of brain structures is highly dependent on the image registration process. The risk of susceptibility from registration between the atlas and the target image, caused by differences in shape, size, or orientation, can introduce errors. Furthermore, a single atlas lacks adaptability to correct for imaging distortions, such as motion artifacts or intensity inhomogeneity, which significantly impact segmentation performance. Another limitation arises from resolution discrepancies. The affine or warping process is prone to under-resampling errors due to the relatively coarse through-plane resolution (3.0–3.5mm) compared to the finer in-plane resolution (0.2– 0.5mm) of NM-MRI or T2*-weighted imaging (Hsu et al., 2025). Additionally, manual tracing remains a highly time-consuming and challenging task due to the low signal-to-noise ratio (SNR) of NM-MRI relative to high-resolution T1-and T2-weighted imaging, raising concerns about reproducibility. In the case of LC segmentation, its location near air-tissue interfaces (e.g., the petrous part of the temporal bone) exacerbates these challenges, as magnetic susceptibility artifacts can distort signals in T2*-weighted imaging, further reducing segmentation accuracy (Haller et al., 2021; Oehler et al., 1995).

Despite being a very small structure, the locus coeruleus (LC) demonstrates noticeable heterogeneity in its cellular composition and circuit organization, with local differences in cell density across subregions (McKinney et al., 2023). Specifically, the LC core contains the highest concentration of cells, whereas the surrounding shell exhibits considerably lower density. Such heterogeneity highlights that the LC is not a uniform structure but rather consists of distinct subregions, some of which appear to serve as primary sites for neuromelanin accumulation (Gilvesy et al., 2022; Veréb et al., 2023). This paper aims to assess the accuracy of automatic LC segmentation using multi-atlas nonlinear warping on high-resolution T1/T2-weighted images, comparing the results with manually defined LC peak regions. Additionally, we analyze T1/T2 intensity within the automatically extracted LC regions to investigate its correlation with NM-MRI, observe age-related trends, and examine T1/T2 intensity changes within the LC region.

## Method

### Participants

This study includes two sets of data. The first dataset was from 134 healthy participants that has 60 male and 74 female, age range between 8 to 64 years who were recruited and T1w/T2w and NM-MRI imaged. All participants provided written informed consent approved by the Virginia Tech Institutional Review Board for examination the accuracy of automatic segmentation of LC and the trend of correlation between NM-MRI and T1/T2 intensity. The second dataset included T1w and T2w images from 652 youth (301 male and 351 female, average age = 14.43, std = 4.07; age range: 5.6 to 21.9), publicly available from Human Connectome Project in Development (HCP-D) (https://www.humanconnectome.org) (Somerville et al., 2018), were used to investigate whether our theories apply in the school-age group. Exclusion criteria included premature birth, and lifetime history of serious medical or endocrine conditions, or their treatment. All participants aged 18 and above provided written informed consent. For children under age 18, a parent or a legal guardian provided informed, written permission for their child to participate in the study.

### Image Acquisition

T1w/T2w structural images and neuromelanin-sensitivity structural images were collected with 3T Siemens MRI scanners, Prisma and Trio, at the FBRI Biomedical Research Institute (FBRI) and the Virginia Tech Corporate Research Center (VTCRC) respectively.

Prisma: The neuromelanin-sensitive images were collected through a T1-weighted FSE imaging sequence (T1-FSE) with the following parameters: repetition time, TR = 750 ms, echo time, TE = 12 ms, flip angle = 120°, two average to enhance the signal-to-noise ratio (SNR), 11 axial slices, field of view (FoV) = 220 mm, bandwidth = 285 Hz/Px, slice thickness = 2.5 mm, slice gap = 1.0 mm, and in-plane resolution = 0.43 × 0.43 mm (Clewett et al., 2016; Sasaki et al., 2006). High-resolution T1/T2-weighted anatomical images were collected with the following parameters: T1w - TR = 2,500 ms, TE = 2.06 ms, flip angle = 8°, bandwidth = 220 Hz/Px, Fov = 256 mm, voxel resolution = 1 mm³ isotropic. T2w - TR = 3,200 ms, TE = 563 ms, flip angle of 120°, bandwidth of 725 Hz/Px, FoV = 256 mm, voxel resolution of 1 mm³ isotropic.

Trio: The neuromelanin-sensitive images were collected through a T1-weighted FSE imaging sequence (T1-FSE) with the following parameters: repetition time, TR = 750 ms, echo time, TE = 11 ms, flip angle = 120°, two average to enhance the SNR, 11 axial slices, field of view (FoV) = 220 mm, bandwidth = 287 Hz/Px, slice thickness = 2.5 mm, slice gap = 1.0 mm, and in-plane resolution = 0.43 × 0.43 mm(Clewett et al., 2016; Sasaki et al., 2006). High-resolution T1/T2-weighted anatomical images were collected with the following parameters: T1w - TR = 2,400 ms, TE = 2.16 ms, flip angle = 8°, bandwidth = 220 Hz/Px, Fov = 256 mm, voxel resolution = 0.8 mm³ isotropic. T2w – TR = 2,500 ms, TE = 497 ms, flip angle of 120°, bandwidth of 574 Hz/Px, FoV = 256 mm, voxel resolution of 1 mm³ isotropic.

### Locus Coeruleus Segmentation

The T1/T2w brain images underwent a correction process for intensity non-uniformity employing the N4 bias field correction algorithm (Tustison et al., 2010), and the corrected images were then rigidly transformed in a standardized stereotaxic space. A brain masking procedure was executed utilizing a majority voting strategy. This involved the joint warping registration of T1w and T2w images with six predefined single or average atlases and the result of the brain extraction tool (BET). Necessary manual corrections to the brain masks were conducted using the itkSNAP software (Yushkevich et al., 2006) to guarantee accurate masks.

The labeling of the locus coeruleus (LC) was defined using five different atlases or multi-modal data to ensure anatomical accuracy and consistency across different reference frameworks. Each atlas provided distinct structural and cytoarchitectural information, enabling a comprehensive delineation of the LC (Edlow et al., 2012; Keren et al., 2009; Tona et al., 2017; Ye et al., 2021). The selection of these atlases was based on their widespread use in neuroanatomical research and their ability to offer high-resolution anatomical details of the brainstem region. By integrating data via majority voting scheme, a more precise and standardized localization of the LC was achieved, minimizing potential discrepancies due to individual atlas variations. This approach facilitated a robust and reproducible method for identifying the LC in neuroanatomical and functional studies (Figure 1). A multi-modality (T1w and T2w) multi-atlas segmentation workflow was employed, utilizing the in-house, open-source MultiSegPipeline software (Cherel et al., 2015). To maintain congruency between the manually delineated LC masks and the automated segmentation outputs, and to rectify any discrepancies in spatial alignment or minor sampling issue from different resolution, an in-plane 2D morphological operation of one voxel dilation followed by erosion was performed. An extensive visual assessment was conducted to ascertain the segmentation quality of all images in terms of anatomical accuracy, leading to the conclusion that no processed data warranted exclusion based on the segmentation quality.

**Figure 1.**
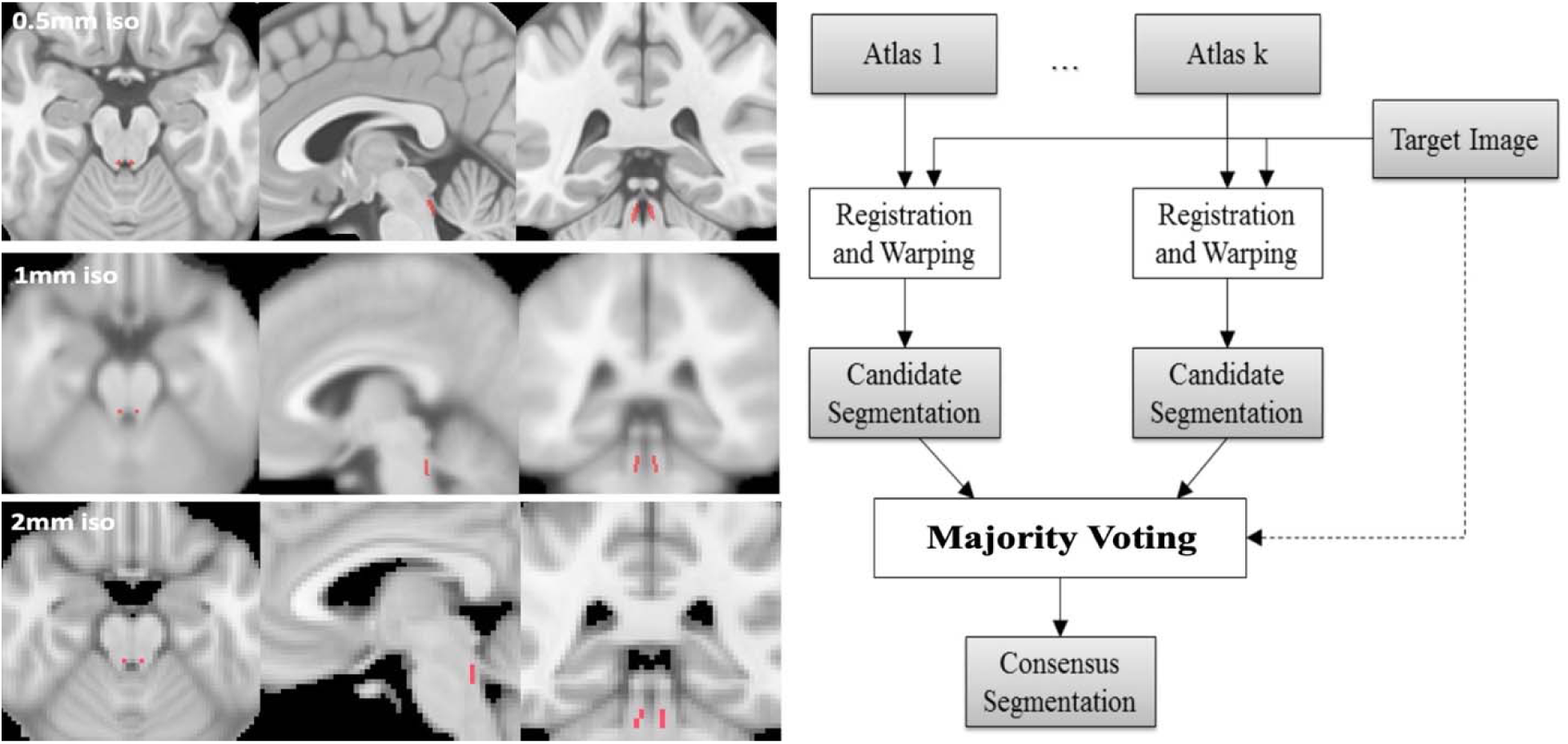
overview. Definition of the locus coeruleus (LC) using five different atlases and multi-modal data to ensure anatomical accuracy and consistency across reference frameworks. Each atlas contributed unique structural or cytoarchitectural details, enabling a comprehensive LC delineation. A majority voting scheme was applied to integrate the atlases, minimizing discrepancies and yielding a precise, standardized LC localization for robust and reproducible neuroanatomical analysis.

The predominant neuromelanin contrast was manually defined following established methods (Clewett et al., 2016). In the axial slice, the inferior colliculus was first identified. Approximately two slices below, the left and right locus coeruleus (LC) regions were located. A 3×3 voxel cross was centered around the peak intensity voxel in each LC hemisphere. As a reference region for calculating the contrast ratio, a 10×10 voxel sample was taken from the dorsal pontine tegmentum (PT), positioned six voxels anterior to the LC crosses and equidistant between them. Given the inherent biases and noise present in NM-MRI or T1/T2-weighted MR images, this reference region was defined to calibrate intensity and ensure accurate contrast measurement.

## Statistical Analysis

In the NM-MNI domain, the overlap between automatically extracted LC labels and manually drawn LC labels were quantified through the counting voxel the overlapping between each other’s. However, all highest intensity measurements were conducted in the each original T1w/T2w or NM-MRI raw space, without applying bias correction or resampling. This approach was chosen to prevent any potential blurring or reduction in intensity values that could result from correction or resampling processes.

Since intensity distribution from extracted LC agreed to the normality, we applied the 2-standard deviation (SD) estimated value that is represented 95% (high/low) value of current distribution. To analyze the normalized LC intensity changes with age, we applied the general linear model built in *Matlab 2024b* using the robust *fitlm* package with a sex as a covariate.

Additionally, the HCP-D data age range from 6 to 22 years was divided into 5 stages (childhood; 6∼9 year, late childhood; 10∼12 year, early adolescence; 13∼15 year, middle adolescence; 16∼17 year, emerging adulthood; 18∼ year) based on developmental milestones, including brain maturation, psychosocial changes, and cognitive growth (Arnett et al., 2000; Casey et al., 2005; Choudhury et al., 2006; Galvan et al., 2006; Luna et al., 2010; Sowell et al., 2003). To assess the age-group differences, a non-parametric permutation test (10,000 iterations) was conducted for each comparison.

## Result

### LC Segmentation and Validation

The automatically extracted locus coeruleus (LC) was overlaid on neuromelanin-sensitive MRI (NM-MRI) in axial (Figure 2a), sagittal (Figure 2b), and coronal (Figure 2c) views. The extracted LC regions were located bilaterally near the fourth ventricle, specifically at the base of its floor, and corresponded closely with manually cross-marked LC areas (blue: overlapping voxels; red: non-overlapping voxels; Figure 2d). A voxel-wise comparison showed that 89.73% of the automatically extracted LC regions shared at least one-voxel with the manual tracings. Applying a 2D in-plane dilation of one voxel (0.25 mm) increased the overlap to 94.52% (Figure 2e). Further dilation to two-voxels (0.5 mm) resulted in a 95.21% match, although this introduced some overestimation into the floor of the fourth ventricle (Figure 2f). When evaluating the coverage of manual tracings, 36.98% of regions achieved perfect overlap (defined as complete coverage of 10 manually marked voxels in both side) and 78.08% achieved partial bilateral overlap (at least one voxel on each left/right side) after one-voxel dilation. These percentages increased to 71.23% and 91.09%, respectively, after two-voxel dilation (Figure 2g).

**Figure 2.**
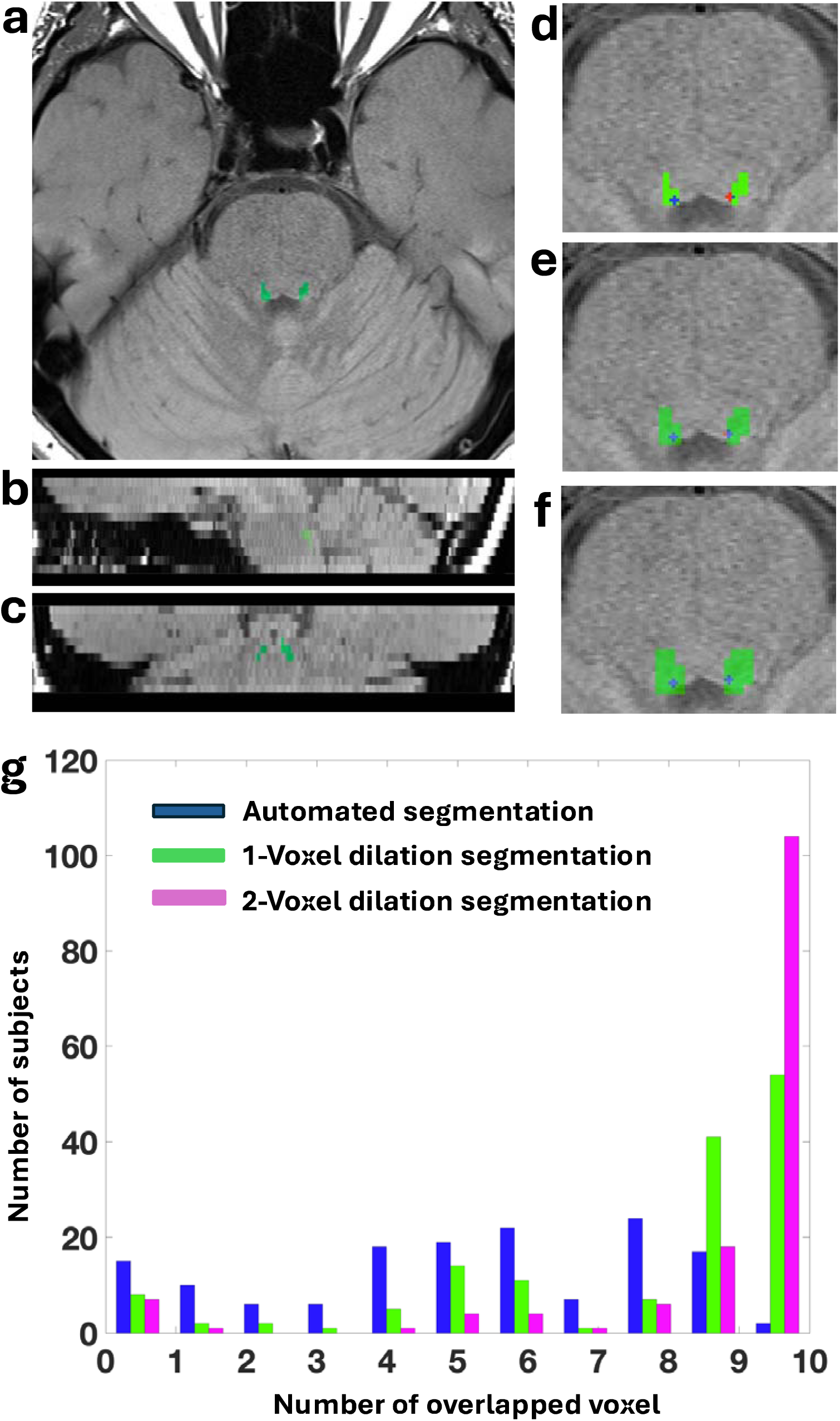
Locus coeruleus segmentation result. The automatically extracted LC ROI labels from T1w/T2w in stereotaxic spaces were transformed into the NM-MRI space and overlaid for comparison (transverse (a), coronal (b), and sagittal (c) views). The overlap between the automatically extracted LC regions without any morphological operations and the manually defined 10-voxel LC masks is shown in (d), with overlapping voxels displayed in blue and non-overlapping voxels in red. Results after one-voxel dilation (e) and two-voxel dilation (f) of the automatically extracted LC regions are also shown. Panel (g) presents the overlap results for all subjects, ranging from cases with no overlap to cases where all 10 voxels overlapped completely.

Figure 3 illustrates the relationship between manually delineated LC intensity and three different intensity measures extracted from automatically segmented LC regions: average intensity (green), statistically estimated peak intensity, defined as the mean plus two times the standard deviation (mean ± 2σ) (blue), and local peak intensity (red). All three measures exhibit a positive linear relationship with the intensity from manually drawn LC, but their performance characteristics differ substantially.

**Figure 3.**
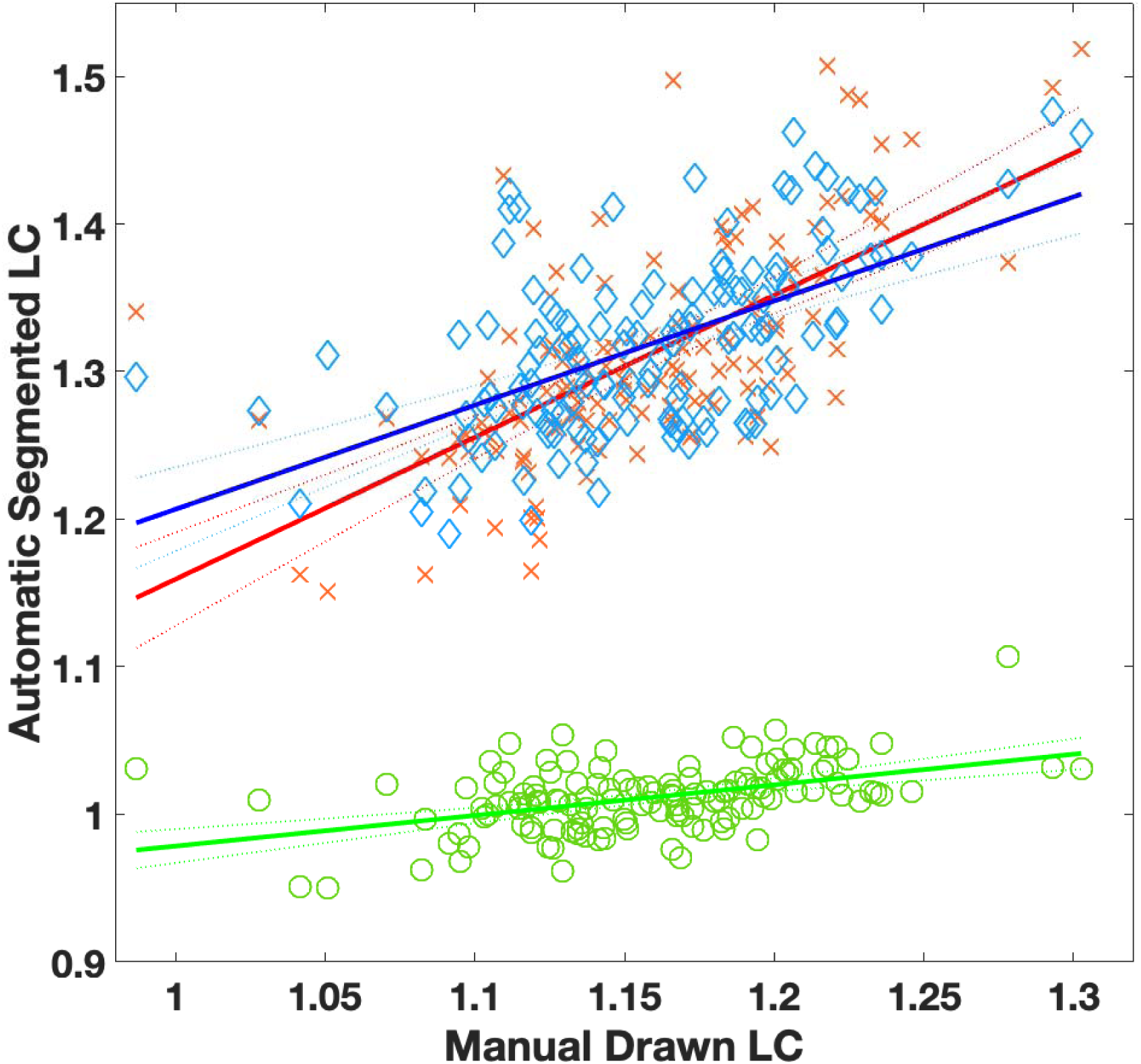
Compare with the average intensity of manually drawn LC and average intensity (green circle, ρ=0.2221(P), ρ=0.2221(S)), maximum (red cross. ρ=0.3499(P), ρ=0.6732(S)) and statistically 95% high estimated intensity (blue diamond, ρ=0.5946(P), ρ=0.5503(S)) of the automatic segmented LC in NM-MRI using Pearson(P) and Spearman(S) correlation.

The average intensity from the automatically segmented LC region demonstrated the relatively low correlation (green circles and line; *Coefficient* = 0.458, *SE* = 0.035). This is likely due to substantial intensity variability within the segmented region—particularly in the LC tail—where inclusion of low-intensity voxels reduces the overall contrast, thereby diminishing correspondence with the real peak values. In contrast, the statistically estimated high intensity across the segmented LC region, showed a stronger linear relationship with the manual LC peak (blue diamonds and line; *Coefficient* = 0.577, *SE* = 0.087). The highest correspondence was observed when using the maximum local peak intensity extracted from the segmented LC region (red crosses and line; *Coefficient* = 0.654, *SE* = 0.097). While this method best reflects the true LC peak intensity, it also yielded the largest standard error among the three, likely due to increased susceptibility to detecting isolated high-intensity voxels caused by noise or segmentation artifacts. Although local peak extraction provides a closer estimate of true LC intensity, statistical estimation methods offer a more reliable and robust measure due to reduced sensitivity to noise.

### Image Processing Effects on LC Intensity and Age Association

MRI intensity varies not only by sequence type such as NM-MRI (Figure 4, A), T1w (Figure 4, B) or T2w (Figure 4, C), but also by intensity inhomogeneity and image processing such as interpolation and resampling. Intensity inhomogeneity in MRI, also known as bias field or non-uniform intensity artifact, is caused by several physical and technical factors that result in spatially varying signal intensity across the image, even when the underlying tissue is homogeneous. N4 Bias Field Correction is a method used to correct intensity inhomogeneity (bias field) in MRI images (Figure 4, D and E). It improves image uniformity by removing smooth, low-frequency variations in intensity caused by scanner imperfections. This correction enhances tissue contrast and improves the accuracy of further analyses like segmentation (Tustison et al., 2010). Rigid transformation to stereotaxic space is done to align each subject’s brain into a common orientation and position without changing its size or shape. This allows for consistent comparison across individuals and compatibility with brain atlases. After applying the six-parameter transformation, the image is resampled in template space (Figure 4, F and G). As a result of the resampling, intensity contrast appears smoother compared to the raw image. Nevertheless, the locus coeruleus exhibits a largely consistent distribution throughout the entire structure. After bias field correction, the LC intensity contrast relative to the reference region changed by an average of 0.285% in the T1-weighted images and 0.289% in the T2-weighted images. In comparison, rigid transformation resulted in larger intensity changes, with mean changes of 0.997% for T1-weighted images and 2.547% for T2-weighted images in our dataset.

**Figure 4.**
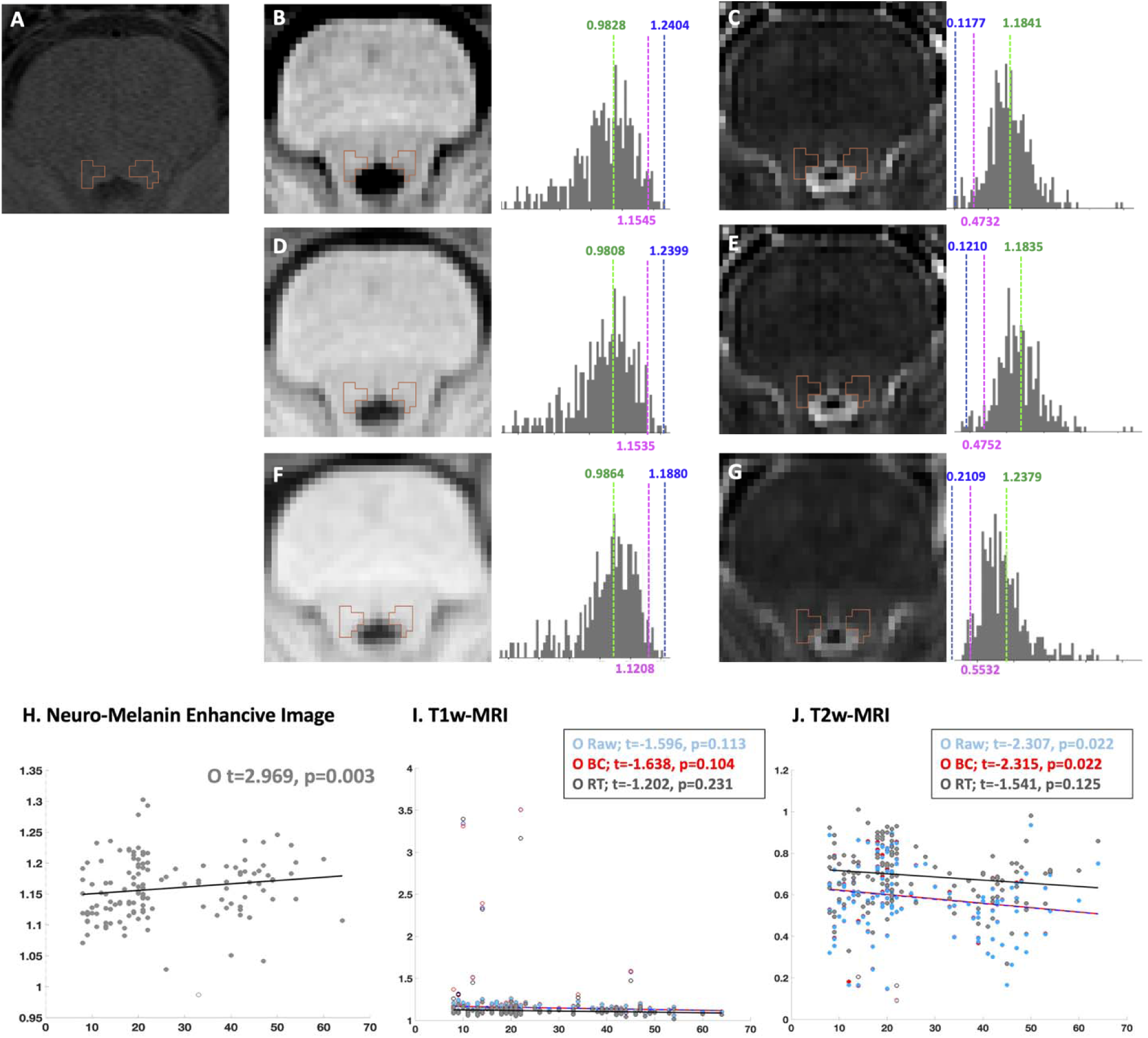
MRI intensity varies by sequence type - A (NM-MRI), B (T1-weighted), and C (T2-weighted) - and is influenced by intensity inhomogeneity (bias field) and image processing steps such as interpolation and resampling. N4 bias field correction improves image uniformity by compensating for low-frequency intensity variations (D, E). Rigid transformation aligns images to stereotaxic space and adjusts intensity contrast through interpolation (F, G). Despite these processing steps, the overall distribution of the locus coeruleus remains consistent across images. For an example subject, the mean intensity (green), mean±2σ (magenta), and mean±3σ (blue) are shown, illustrating intensity changes after processing. Notably, mean±3σ represents extreme values covering approximately 99.5% of the data, but may also reflect outliers that fall outside the actual data distribution. Neuromelanin-sensitive images show a significant positive correlation between LC intensity and age (H). In contrast, T1-weighted images show no significant association with age (I), and T2-weighted images display a significant age-related pattern consistent with NM-MRI (J).

With increasing age, neuromelanin progressively accumulates in the LC, resulting in a statistically significant positive correlation in neuromelanin-sensitive images (Figure 4, H). However, intensity values extracted from T1-weighted images did not exhibit a statistically significant association with age (Figure 4, I). Notably, after removing approximately 6% of data points identified as potential outliers (e.g. due to motion artifact, magnetic susceptibility, error of bias field correction or reconstruction artifact) the correlation not only remained non-significant but also shifted toward a negative direction, contrary to our initial hypothesis (S.Figure 2). In contrast, intensity measures derived from T2-weighted images showed a statistically significant pattern consistent with age-related neuromelanin accumulation, even though the statistical power tended to diminish slightly following rigid transformation (Figure 4, J).

### Developmental Emergence of Detectable Neuromelanin in the LC Across Childhood to Early Adulthood

Neuromelanin generally becomes detectable only after a certain level of accumulation, making it more readily observable in the brains of older individuals compared to younger ones. Therefore, we utilized the HCP-D dataset to investigate the developmental trajectory of neuromelanin in a younger population. While LC intensity extracted from T1-weighted images did not show a clear age-related trend (Figure 5, A), intensity values derived from T2-weighted images exhibited a statistically significant negative regression across the 6-to 22-year age range (Figure 5, B). This finding is consistent with previous results obtained from neuromelanin-sensitive imaging and T2-weighted data To determine the age at which accumulated neuromelanin becomes detectable, we divided the sample into five developmental age groups and examined differences in mean LC intensity contrast across age groups. Figure 5, C and D show that no significant differences were observed in either T1w or T2w images during childhood and early adolescence (group 1∼3). However, beginning in middle adolescence (group 4, age 16∼17), differences in LC intensity contrast began to emerge. For LC intensity contrast extracted from T1w image, Groups 4 and 5 showed a modest but statistically significant difference compared to childhood (group 1, age 6∼9) (p=0.0095 and p=0.0424). However, no significant differences were observed between adjacent developmental groups. Notably, LC intensity measured from T2w images showed a statistically robust difference in the late adolescence or emerging adulthood group (group 5, after age 18) compared to all other age groups (group 2-5; p=0.002, group 3-5; p=0.0015 and group 4-5; p<0.0001).

**Figure 5.**
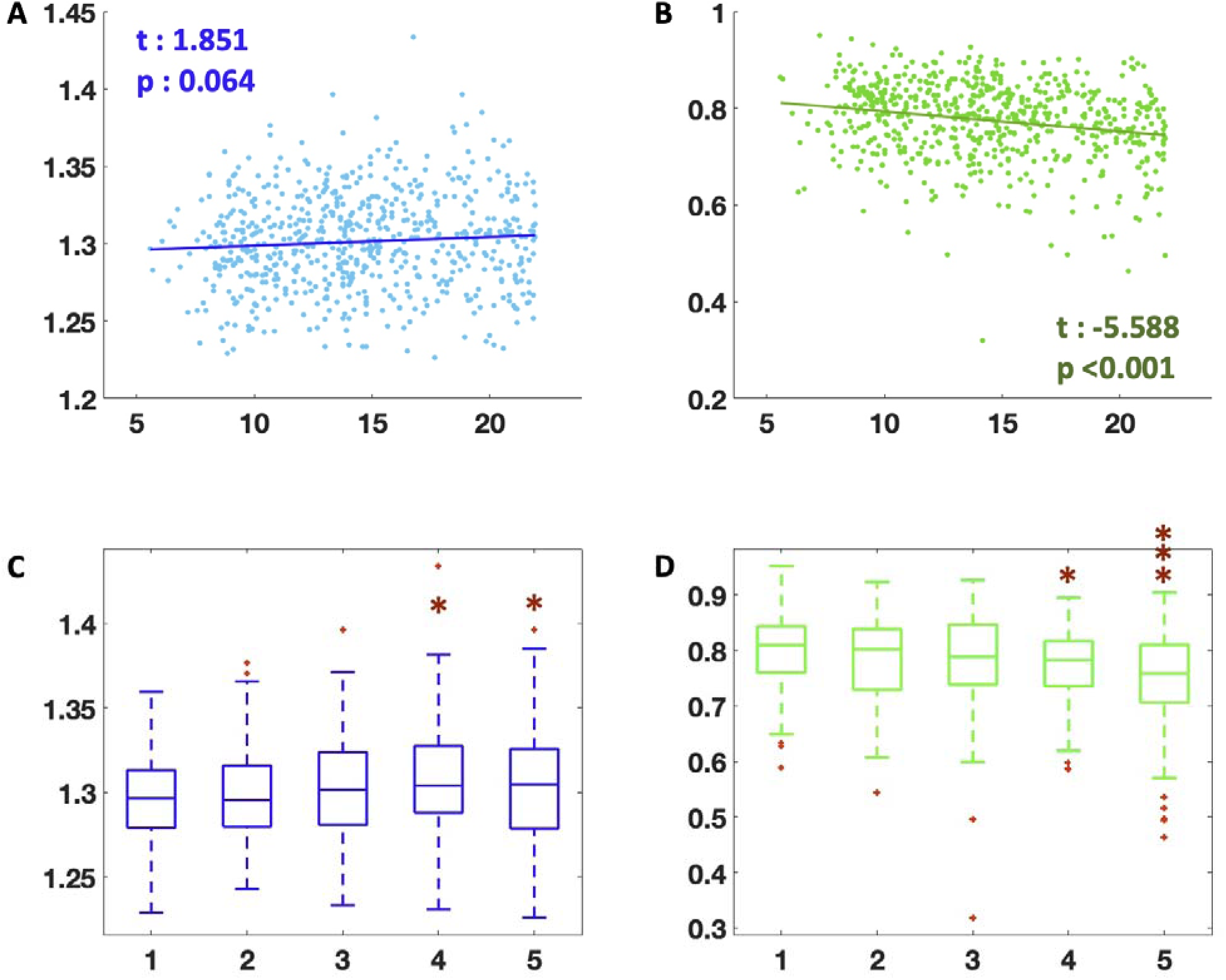
Developmental alteration of LC intensity across childhood to early adulthood. (A) LC intensity contrast extracted from T1-weighted images shows no significant association with age (t = 1.851, p = 0.064). (B) In contrast, LC intensity contrast from T2-weighted images shows a significant negative association with age (t =-5.588, p < 0.001), indicating decreasing intensity across the age range of 6 to 22 years. (C) Boxplots of T1-weighted LC intensity changes across five developmental age groups (1: age 6∼9, 2: age 10∼12, 3: age 13∼15, 4: age 16∼17, 5: age ≥18) show no significant differences among early groups, but groups 4 and 5 show modest increases compared to group 1. (D) Boxplots of T2-weighted LC intensity changes show a statistically significant decrease in group 5 (late adolescence/emerging adulthood) compared to all younger groups.

## Discussion

### Automated LC segmentation

Automated segmentation of the locus coeruleus (LC) has traditionally relied on prior information or probabilistic maps derived from population-specific templates. However, the use of single-source priors can introduce systematic biases and may not generalize well across diverse populations or imaging protocols. As a result, there is a growing interest in multi-modal and multi-atlas frameworks to enhance robustness and accuracy.

Recent advancements have also demonstrated the utility of deep learning-based models, which leverage data-driven training to achieve more refined and reliable LC segmentation (Dünnwald et al., 2021). Although various approaches have been developed to improve the accuracy of locus coeruleus (LC) segmentation, current methods typically achieve only 50∼70% accuracy (Ariz et al., 2019; Dünnwald et al., 2021; Tona et al., 2019). This limitation arises from the LC’s inherently small, slender structure and its low intensity contrast relative to adjacent brainstem tissue, which complicates reliable delineation. The challenge is particularly pronounced in the caudal portion of the LC, where anatomical variability and reduced signal conspicuity further hinder precise segmentation.

Moreover, within the LC region, the degree of high-intensity signal reflecting neuromelanin accumulation serves as a critical biomarker, given its established relevance to both clinical phenotypes and underlying neurobiological processes (Clewett et al., 2016; Li et al., 2022; Liu et al., 2019). Therefore, it is critical to evaluate how accurately the automatically segmented LC region encompasses the peak intensity areas, as these regions are key to neuromelanin-based biomarker analyses. While our study employed 3D volumetric segmentation to enable furthermore investigations such as volumetric quantification, we emphasized the importance of validating this segmentation against manually delineated peak intensity regions. As demonstrated in our results, applying the morphological operation of a one-voxel dilation enables the inclusion of most peak intensity voxels within the segmented LC mask. This suggests that the proposed automated method can reliably capture the brightest (or darkest) regions in each individual images, providing a stable foundation for intensity-based analyses.

For the final label generation, label fusion methods that incorporate local intensity similarity between atlases and the target image, such as joint label fusion, are commonly employed in complicated structures requiring high segmentation accuracy (Wang et al., 2013). However, these methods are computationally demanding and require careful tuning of parameters, such as patch size. In contrast, the locus coeruleus (LC) exhibits a relatively simple structure with minimal inter-atlas variability, allowing simple majority voting to achieve sufficiently accurate segmentations in this region.

### Average or estimated peak intensity in NM-MRI

In neuroimaging studies, the mean or median is commonly used as a representative value for a given region, as it effectively reflects the central tendency of the distribution by incorporating all voxel intensities. This approach provides a robust summary of the overall characteristics of the region (Giorgi et al., 2022; Liu et al., 2020; Liu et al., 2017; Tona et al., 2019). However, in the LC, where neuromelanin accumulation tends to concentrate in the head region and diminishes toward the tail portion, peak intensity values have often been employed to capture these localized high-intensity signals (Berger et al., 2023; Dahl et al., 2019; Liu et al., 2025). Despite its intuitive appeal, the use of peak intensity in neuroimaging carries substantial limitations. Peak values are highly susceptible to random noise, increasing the risk of extracting unreliable or spurious measurements. Moreover, even within the same participant and scanning protocol, the exact location of the peak intensity can vary across sessions, reducing its reliability for longitudinal or group-level comparisons. Peak-based measures inherently reflect extreme values rather than central tendencies, resulting in greater within-group variability compared to metrics such as the mean or median, which can subsequently increase the risk of type I errors in statistical analyses (Lieberman & Cunningham, 2009; Schwartzman et al., 2011).

To address the challenges associated with extreme outliers, Riley et al. employed an intensity rescaling procedure in their study, in which turbo spin echo images were normalized within each region of interest (ROI) using a ±3 standard deviation range, encompassing 99.7% of the data under the assumption of a normal distribution. This approach was selected to approximate min–max normalization (scaling intensities between 0 and 1) while simultaneously mitigating the influence of noise-driven extreme outliers. By excluding extreme voxel intensities that may arise from noise or artifacts, this rescaling procedure offers a practical compromise between data normalization and robustness (Riley et al., 2025). However, while this approach can effectively suppress the influence of extreme outliers, it may still retain extreme values, as the ±3 standard deviation range can, in some cases, encompass such extreme intensities.

We derived a representative high-intensity value as the mean ± 2σ (which represented 95.4% of the data) from the LC intensity distribution after normalizing by the average intensity of the reference region, pontine tegmentum, as a robust and interpretable summary measure. In figure 3, both the statistically estimated peak value (mean ± 2σ) and the extracted local peak intensity showed stronger associations with the manually defined LC intensity. Notably, while the local peak (from automatic segmented LC ROI) provided the closest approximation to the true LC peak in NM-MRI, it also showed a larger standard error-likely reflecting sensitivity to local noise or spurious high-intensity voxels within the segmented region. On the other hand, the statistical estimation method offered a more stable and noise-resilient alternative, effectively capturing the signal characteristics of the high-intensity LC core. These results suggest that for robust LC signal quantification, especially in large-scale or noisy datasets, statistical peak estimation may offer a more reliable surrogate marker than either the average intensity or raw local peak.

### Locus Coeruleus’ intensity changes on T1w and T2w image

During neurotransmission, norepinephrine undergoes enzymatic degradation, primarily by monoamine oxidase (MAO) (Edmondson, 1995; Pástorová & Várady, 1996). This process produces various reactive byproducts, such as catechol-aldehydes and other oxidized metabolites, which can be potentially cytotoxic due to their capacity to induce oxidative stress. These reactive byproducts are further oxidized and polymerized into neuromelanin, a dark, insoluble pigment composed of complex polymers bound to lipids and metal ions (e.g., iron, copper). This process gradually converts potentially harmful substances into an inert form, leading to their long-term sequestration within neurons. Neuromelanin accumulates slowly within lysosome-like structures inside LC neurons. The amount of neuromelanin increases with age, making it a prominent feature in adult and elderly brains, while it is barely detectable in childhood (Capucciati et al., 2021; Halliday et al., 2006; Zucca et al., 2018).

Given the variety of imaging modalities available for the locus coeruleus (LC), it is important to summarize their key characteristics and relative advantages. T1-weighted imaging highlights tissues based on differences in longitudinal relaxation time. Tissues with high lipid content or paramagnetic substances shorten T1 and appear bright (Paus et al., 2001). The LC is difficult to detect on standard T1w imaging due to limited contrast against surrounding brainstem tissue. Therefore, the accumulated intensity within the LC does not seem to exhibit clear age-related variations (Figure 4, I and Figure 5, A). However, the LC contains neuromelanin, a pigment with paramagnetic properties that can enhance T1 contrast. Therefore, Specialized T1w techniques, such as neuromelanin-sensitive MRI, can enhance the visibility of the LC by exploiting its neuromelanin content. This technique uses magnetization transfer effects to improve contrast. The neuromelanin-sensitive MRI, typically based on T1-weighted sequences, is considered the most effective technique for visualizing LC structure (Sasaki et al., 2008). This approach takes advantage of the intrinsic contrast provided by neuromelanin, which accumulates within LC neurons over time, allowing for clear delineation of the LC. Although neuromelanin-sensitive MRI (NM-MRI) is widely used for visualizing the locus coeruleus (LC), it has several limitations. Its specificity is limited, as signal intensity can be influenced by factors beyond neuromelanin, such as water content and macromolecular structures. The LC is also prone to motion and susceptibility artifacts, which may affect image quality. Furthermore, NM-MRI relies on relative contrast rather than direct quantification, and its sensitivity is reduced in younger individuals due to low neuromelanin levels. Finally, NM-MRI results can vary depending on scanner type, field strength, and imaging protocols, limiting cross-study comparability (Hwang et al., 2023; Priovoulos et al., 2020).

On the other hand, T2-weighted imaging is sensitive to differences in transverse relaxation time, often highlighting fluid-rich areas where water content is higher. The LC is poorly visualized on standard T2w MRI. The signal contrast in T2w images is generally low in the brainstem and does not effectively differentiate the LC from surrounding white matter. However, the LC may have some iron deposition that alters the local magnetic environment by increasing magnetic susceptibility effects within the LC. These susceptibility effects lead to a shortening of T2 relaxation times, causing a decrease in signal intensity on T2-weighted images. Consequently, the LC appears increasingly hypointense (darker) with advancing age (Figure 4, J and Figure 5, B). In contrast, T2-weighted imaging techniques, such as susceptibility-weighted imaging (SWI) or quantitative susceptibility mapping (QSM), are more sensitive to iron deposition and microvascular properties (Betts et al., 2019; Hashido & Saito, 2016; Trujillo et al., 2019; Zang et al., 2025). These methods are less specific to the LC itself, as their signal changes may reflect a variety of sources beyond neuromelanin content.

Overall, conventional T1-weighted imaging has limited sensitivity for directly or indirectly detecting neuromelanin in the LC. In contrast, conventional T2-weighted imaging may allow for the indirect assessment of age-related neuromelanin accumulation through signal reductions, similar to the changes observed in NM-MRI.

### Locus Coeruleus’ intensity changes in the young age group

Neuromelanin accumulation in the human brain begins at a relatively late developmental stage compared to other neurobiological processes. During early childhood, neuromelanin is either absent or present at minimal levels, making it virtually undetectable using conventional imaging techniques. Several postmortem and neuroimaging studies indicate that neuromelanin accumulation in the locus coeruleus (LC) and substantia nigra (SN) begins during adolescence, typically around the onset of puberty (Beardmore et al., 2021; Zucca et al., 2006).

Zucca et al invested that neuromelanin levels remain low but gradually start to increase in the early adolescent period (approximately 13∼15 years). Noticeable accumulation becomes more apparent during mid to late adolescence (approximately 16∼20 years), with further increases occurring in young adulthood (Zucca et al., 2006). Our findings from the Human Connectome Project Development dataset are consistent with previous reports. We observed only subtle increases in neuromelanin-related signal intensity during childhood and early adolescence (approximately ages 13 to 15), with changes that were too small to reach statistical significance. However, from around age 16, corresponding to middle and late adolescence, we detected statistically significant changes—either increases or decreases—in LC intensity signals associated with neuromelanin accumulation (Figure 5, C and D). Throughout adulthood, neuromelanin continues to accumulate progressively, with peak levels generally observed in middle to late adulthood (typically after the age of 50–60). Notably, neuromelanin accumulation shows a strong positive correlation with age in adults, making it a potential biomarker for age-related brain changes.

## Conclusion

This study highlights several key considerations in the segmentation and quantification of the locus coeruleus (LC). While recent advances in automated LC segmentation, including approaches based on deep learning, have improved reliability, current methods still face challenges due to the LC’s small size, low contrast, and anatomical variability. Importantly, our findings emphasize the need to evaluate whether automated segmentation adequately captures high-intensity neuromelanin-rich regions, as these areas are critical for biomarker analyses. In terms of LC intensity quantification, we found that statistically estimated high-intensity values (mean ± 2σ) offer a more stable and noise-resilient measure compared to local peak intensity, particularly in large-scale or noisy datasets. This approach may serve as a robust surrogate marker for neuromelanin-related signal in the LC.

Our analyses also provide insight into age-related changes in LC signal across different imaging modalities. Conventional T1-weighted imaging showed limited sensitivity for detecting neuromelanin accumulation, while T2-weighted imaging revealed age-related signal reductions likely reflecting neuromelanin and iron accumulation, similar to patterns observed in neuromelanin-sensitive MRI (NM-MRI).

Finally, our developmental trajectory analysis confirms that neuromelanin accumulation in the LC begins gradually during adolescence, with statistically significant changes emerging around mid-to-late adolescence (ages 16 and older). These findings support the notion that neuromelanin-sensitive MRI or T2w image is more suited for adolescent and adult populations, and that careful age consideration is necessary in LC imaging studies. Together, our findings provide practical methodological recommendations for LC segmentation and signal quantification and offer new insights into age-dependent changes in LC imaging.

## Funding

Grants and funding: This work was done based on research supported by R01AG075000/AG/NIA NIH and by 2022 Virginia Tech Lay Nam Chang Dean’s Discovery Fund.

## Author Contributions

SHK: Conceptualization, Supervision, Investigation, Methodology, Formal analysis, Writing; MAS: Supervision, Writing; THL: Conceptualization, Methodology, Writing; JN, & YYC: Data collection and preparation.

## Notes

### Competing Interest Statement

The authors have declared no competing interest.

