## Supplemental Figure1 for "Automated Locus Coeruleus segmentation and age-related signal changes across Neuromelanin-sensitive MRI and T1/T2-weighted images"

### Supplementary

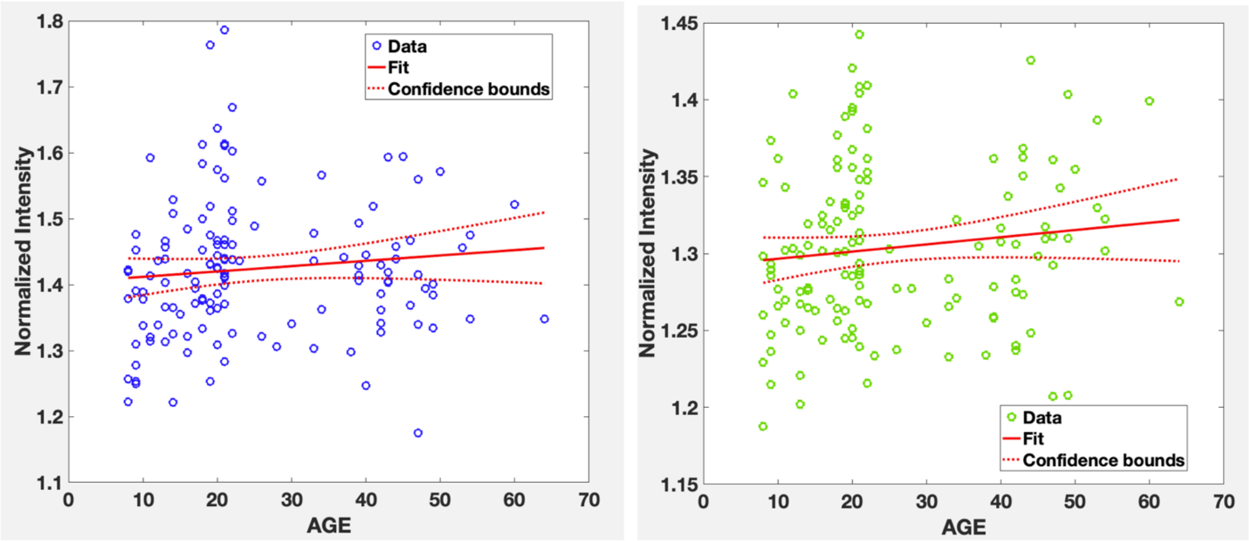


Manual Segmented LC          Automatic Segmented LC

*S.Figure1. The average NM-MRI intensity values of the manually drawn LC (indicated by the blue circle on the left) and the automatically segmented LC (indicated by the green mark on the right) are shown. Since the manual LC was marked only at the peak region, it maintains a higher average intensity compared to the automatically segmented method, which captures the entire LC body. However, both methods show a similar pattern of NM-MRI intensity increase with age.*
