## Supplemental Figure2 for "Automated Locus Coeruleus segmentation and age-related signal changes across Neuromelanin-sensitive MRI and T1/T2-weighted images"

**Supplementary**

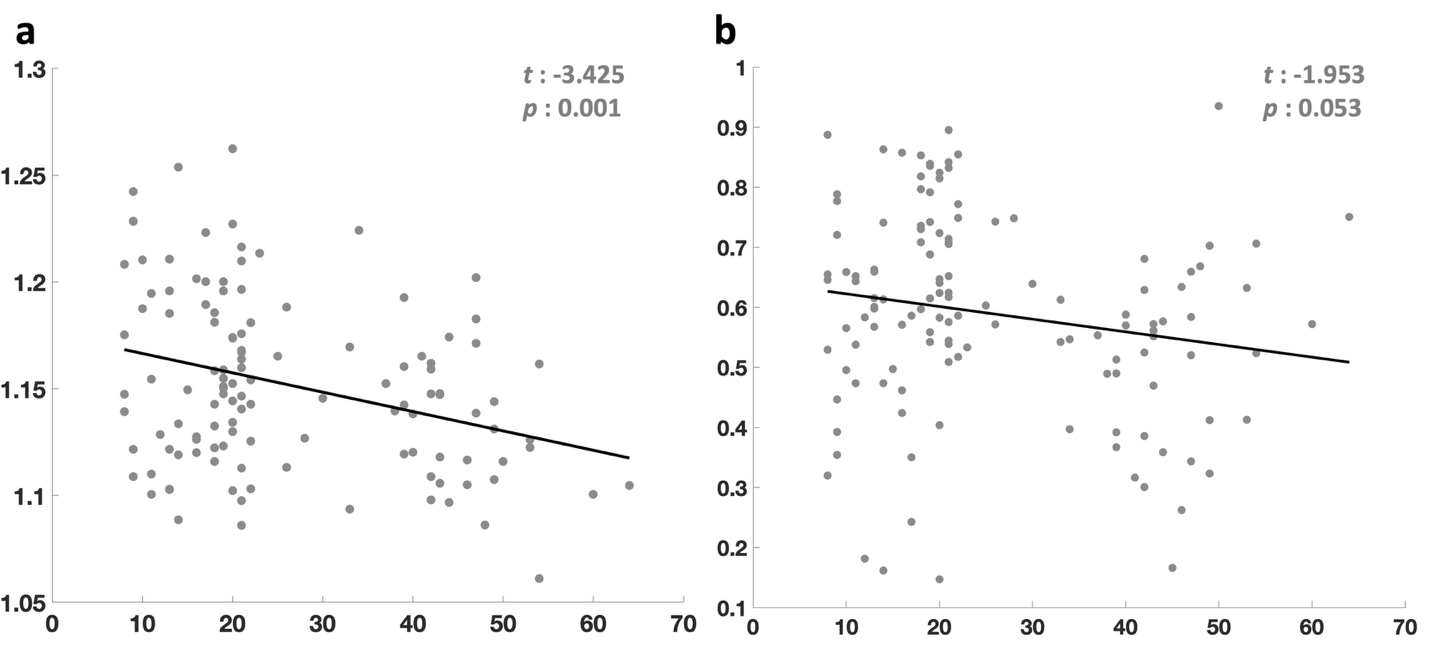

S.Figure 2. This plot shows the data after removing outliers from the raw T1w/T2w values. In figure a, removing 6.2% of outlier data from the T1w image reveals a statistically significant negative correlation with age. In figure b, after excluding 0.8% of outlier data from the T2w image, the statistical power slightly decreases, but a meaningful negative relationship with age remains.
