## Supplemental Figure3 for "Automated Locus Coeruleus segmentation and age-related signal changes across Neuromelanin-sensitive MRI and T1/T2-weighted images"

### Supplementary

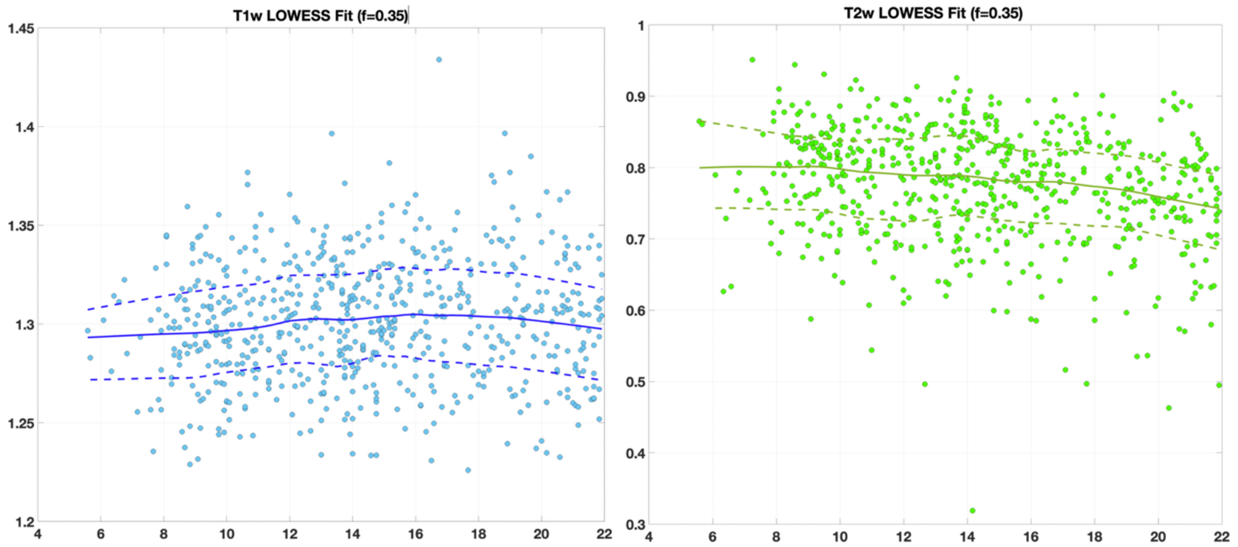


*S.Figure 3. Scatter plots showing the relationship between age and locus coeruleus (LC) intensity contrast in T1-weighted (T1w; left) and T2-weighted (T2w; right) MRI images. Each point represents an individual participant in HCP-D data set. The solid lines indicate locally weighted scatterplot smoothing (LOWESS) fits (f = 0.35), with dashed lines showing the associated confidence bands. LC intensity contrast from T1w images shows minimal variation across ages, whereas T2w images exhibit a slight negative trend, suggesting a modest decline in LC contrast with increasing age.*
